# Homocysteine accelerates hepatocyte autophagy by upregulation of TFEB via DNMT3b-mediated DNA hypomethylation

**DOI:** 10.1101/2023.01.30.526165

**Authors:** Anning Yang, Wen Zeng, Yinju Hao, Hongwen Zhang, Qingqing Wang, Yue Sun, Shangkun Quan, Ning Ding, Xiaoling Yang, Jianmin Sun, Huiping Zhang, Bin Liu, Yun Jiao, Kai Wu, Yideng Jiang

## Abstract

Autophagy plays a critical role in the physiology and pathophysiology of hepatocytes. High levels of homocysteine (Hcy) promote autophagy in hepatocytes, but the underlying mechanism is still unknown. Here, we investigated the relation between Hcy increased autophagy levels and the expression of nuclear transcription factor EB (TFEB). We demonstrate that Hcy increased autophagy levels is mediated by upregulation of TFEB. Silencing TFEB decreases the autophagy-related protein LC3BII/I and increases p62 expression levels in hepatocytes after exposure to Hcy. Moreover, the effect of Hcy on the expression of TFEB is regulated by hypomethylation of TFEB promoter catalyzed by DNA methyltransferase 3b (DNMT3b). In summary, this study shows that Hcy can activate autophagy by inhibiting DNMT3b-mediated DNA methylation and upregulating TFEB expression. These findings provide another new mechanism for Hcy-induced autophagy in hepatocytes.

## INTRODUCTION

Homocysteine (Hcy) is an intermediate of methionine metabolism [1]. Dysregulation of homocysteine metabolism leads to hyperhomocysteinemia (HHcy), an independent risk factor for cardiovascular and cerebrovascular diseases [2]. The liver is the main organ for Hcy-metabolism, however, liver injury, such as chronic liver disease, liver cirrhosis, or even primary liver cancer, the results in an increased level of Hcy in circulation [3-5].

The transcription factor EB (TFEB) is a downstream regulatory factor of rapamycin target protein complex 1 (mTORC1) and is negatively modulated by mTORC1 [6]. It has a vital role in lysosomal biosynthesis, autophagy, and angiogenesis and promotes lipid degradation *in vivo* [7]. Under physiological conditions, TFEB is inactive in the cytosol, whereas fasting and stress stimuli induce its dephosphorylation-mediated activation and subsequent nuclear translocation. Previous studies have shown that TFEB is the primary regulator of autophagy [8]. TFEB actively regulates autophagy, activates lysosomal genes, promotes the formation and fusion of autophagosomes and lysosomes, increases the process of autophagic flux, and enhances the ability of cells to degrade lysosomal substrates [9]. Moreover, TFEB also enhances lipid decomposition and liver lysosomal enzyme activity in fulminant hepatitis mouse model [10]. However, whether TFEB mediates the effect of Hcy on hepatocyte autophagy remains unclear.

DNA methylation, as a type of epigenetic regulation, inhibits DNA methyltransferase mediated gene expression [11, 12]. Hcy is a methyl donor and is involved in gene expression changes by DNA methylation [13]. Meanwhile, HHcy in mammals regulates the expression and activity of DNA methyltransferases (DNMTs) including DNMT1, DNMT3a, and DNMT3b. For instance, cystathionine-beta-synthase deficiency mouse-induced HHcy leads to a significant increase in DNMT activity [14]. Similarly, Hcy promotes DNMT1 protein expression in human umbilical vein endothelial cells [15]. DNMT1 and DNMT3a expression was elevated, while DNMT3b expression was decreased in Hcy-induced mouse brain endothelial cells [16]. However, whether DNA methylation affects the expression of Hcy-induced TFEB and the exact mechanisms have not been fully elucidated.

In this study, we investigated the precise regulatory role and mechanism of TFEB in Hcy-induced autophagy in hepatocytes. Our results revealed that TFEB is an important regulatory mediator of autophagy, and its expression is modulated by DNA methylation. These findings provide novel insights into the molecular mechanism underlying Hcy-induced autophagy in hepatocytes.

## MATERIALS AND METHODS

### Materials

L-Hcy and the DNA methylation inhibitor-5-azacytidine (AZC) were obtained from Sigma-Aldrich (6943, A23885, USA). DC-05, Theaflavin-3, 3′-digallate (TF-3), and Nanaomycin A (NA) were purchased from MedChemExpress (Monmouth, HY-12746, HY-N1992, HY-103397, USA).

### Cell culture

The Human hepatocyte line (HL-7702) was derived from the Cell Bank of the Chinese Academy of Sciences (Shanghai, China) and plated in RPMI-1640 medium (Thermo Fisher Scientific, USA) containing 8% fetal bovine serum (FBS) (HyClone, USA), streptomycin (100 μg/mL) and penicillin (100 U/mL). Then, when cell fusion reached 80% confluence, subsequent experiments were performed after 24 h of intervention with L-Hcy (100 μmol/L).

### Quantitative real-time PCR (qRT-PCR)

qRT-PCR analysis of genes was performed as described previously [17]. Total RNA was extracted using a commercial kit (Takara, Japan) in line with the manufacturer’s protocol. Single-strand cDNA was generated with a SuperScript™ IV One-Step RT-PCR System (ThermoFisher Scientific, USA). qRT-PCR was performed using PowerTrack™SYBR Green Master Mix (Thermo Fisher Scientific, USA) in an ABI7500 (Applied Biosystems, USA). Glyceraldehyde phosphate dehydrogenase (GAPDH) served as internal reference gene. Relative quantification of mRNA expression was calculated using the 2FF0D-^ΔΔCt^ method. The primer sequences for mRNA expression analysis are shown in table 1.

**Table 1.**
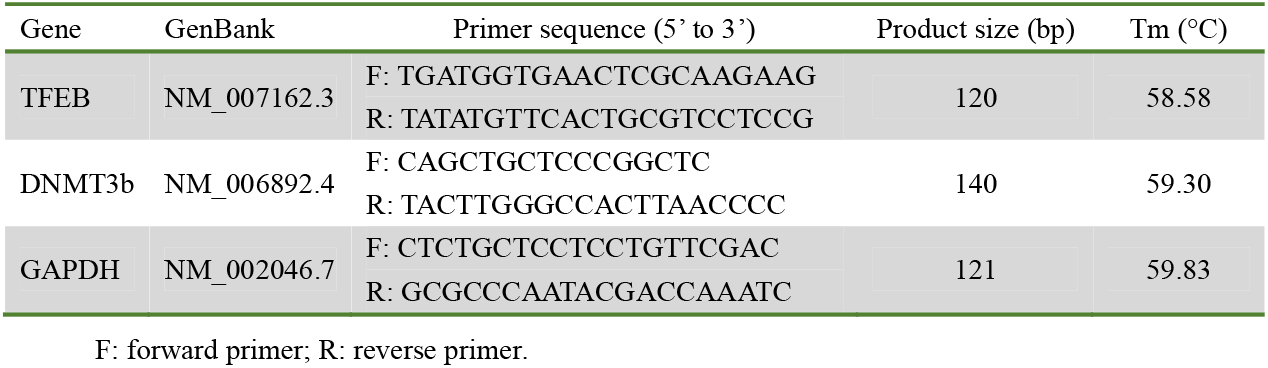
Primer sequence for qRT-PCR analysis.

### Immunoblotting analyses

Hepatocytes were lysed with lysis buffer supplemented with phenyl methane sulfonyl fluoride as previously described [17]. After centrifugation, protein concentrations in the extracts were analysed using the BCA assay (Beyotime Institute of Biotechnology, China). Proteins (30 μg from each extract) were separated by SDS-PAGE gels, and then according to protein molecular weights on gels, extracts were transferred to polyvinylidene fluoride (PVDF) membranes (Merck Millipore, German). Membranes were blocked with 5% defatted milk and then incubated with primary antibodies against anti-DNMT1 (Thermo Fisher Scientific, PA5-30581, USA), anti-TFEB (ab267351), anti-DNMT3a (ab2850), anti-DNMT3b (ab2851), anti-p62 (ab109012), anti-microtubule-associated protein 1 light chain 3 (anti-LC3) (ab192890), and anti-ACTB (ab8226), respectively. Primary antibodies were obtained from Abcam Inc USA. Subsequently, secondary horseradish peroxidase (HRP)-labelled antibodies were used for detection of the indicated protein. Immunoblot images were analysed with ChemiDoc (Bio-Rad, USA).

### Nested methylation-specific-polymerase chain reaction (nMS-PCR)

Genomic DNA was isolated from the hepatocytes using the WizardÒGenomic DNA Purification Kit (Promega, USA). This integrated the DNA denaturation and bisulfite conversion processes into one-step by the EZ DNA Methylation-Gold^™^ Kit (ZYMO Research, USA). The eluted DNA sample was stored at 20°C. The first step of nMS-PCR used an outer primer pair set that did not include any CpG. The second-step PCR was performed with the conventional PCR primers. The sequences of the primers used for the nMS-PCR assays is shown in Table 2. PCR products were purified with an agarose gel. To reduce misprinting and increase efficiency, touchdown (TD) PCR was used for amplification. Samples were subjected to 25 cycles in a TD program (95°C for 30 s, 56.5°C for 30 s, and 72°C for 1 min), followed by a 0.5°C decrease in the annealing temperature every cycle. After completing the TD program, 20 cycles were subsequently run (95°C for 35 s, 50°C for 35 s and 72°C for 35 s), ending with a 3 min extension at 72°C. The PCR products were separated by electrophoresis through a 2% agarose gel containing ethidium bromide. DNA bands were visualized by ultraviolet light. The presence of methylation was calculated using the following formula: Methylation % = Methylation/ (methylation + unmethylation) ×100%.

**Table 2.**
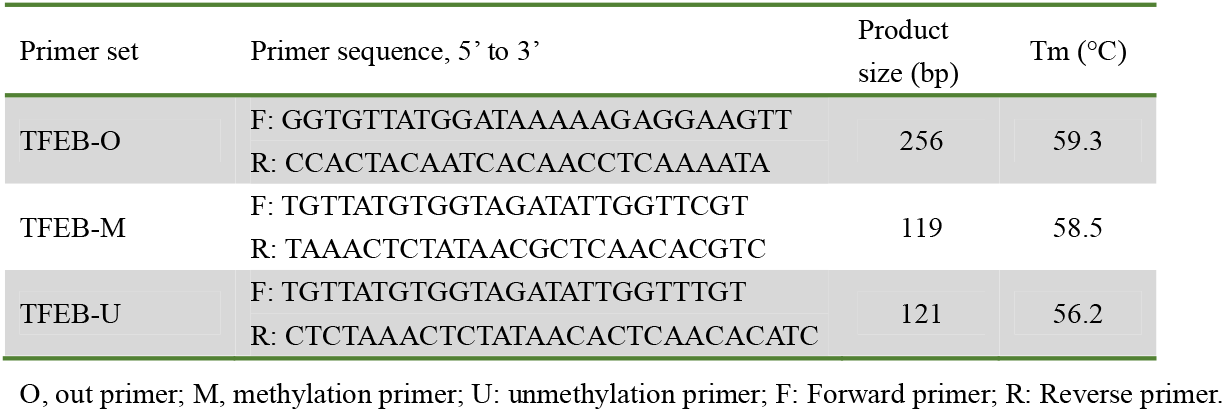
nMS-PCR primers for TFEB.

### Construction of recombinant DNMT3b adenovirus

Recombinant adenoviruses expressing the human DNMT3b gene were constructed with replication-defective adenoviral shuttle vector pHBAd-CMV-IRES-GFP and the adenoviral backbone plasmid pBHGlox(Delta)E1,3Cre. The DNMT3b fragments were inserted into the pHBAd-CMV-IRES-GFP vector and cotransfected with pBHGlox(Delta)E1,3Cre into the virus packaging cell line 293. Recombinant adenoviruses were expanded, purified, collected, and used to infect the liver cell line HL-7702. The recombinant adenovirus encoding green fluorescent protein (Ad-GFP) was used as a control. Hepatocytes at approximately 80% confluence were infected with purified adenovirus for further experiments. Immunoblotting was used to detect ectopic gene expression by antibodies against DNMT3b.

### Construction of TFEB and DNMT3b shRNA adenovirus

For knockdown of TFEB and DNMT3b by short hairpin (shRNA), shRNA adenoviral particles were purchased from Genepharma (Shanghai, China) and packaged into HEK293 cells according to the manufacturer’s guidelines. The sequences of TFEB and DNMT3b shRNA are listed in Table 3.

**Table 3.**
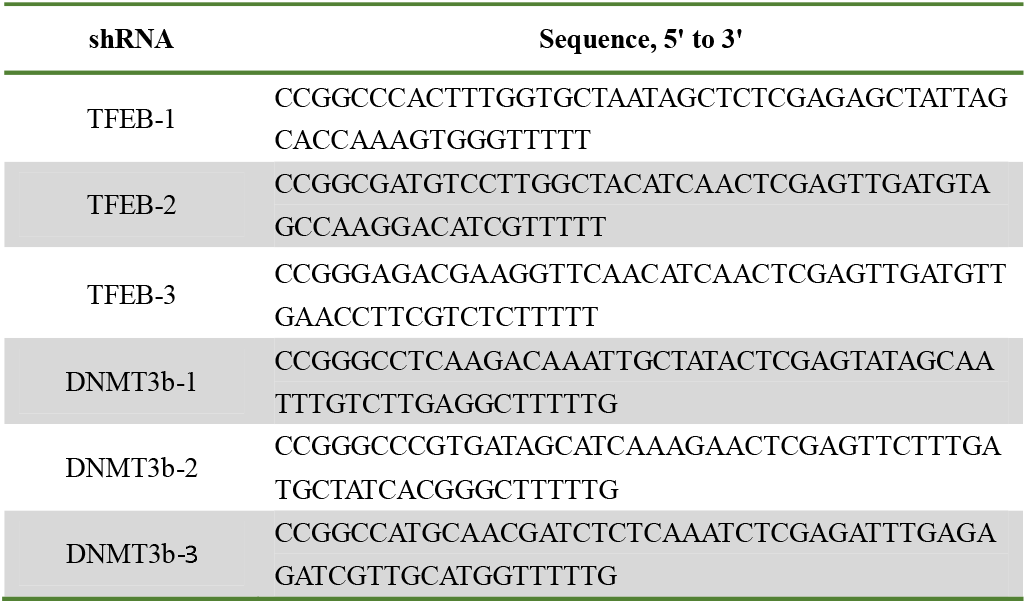
Sequences of shRNA against TFEB and DNMT3b.

### MassARRAY DNA methylation analysis

The Agena MassARRAY platform (Biomiao Biological, China) was used for the quantitative analysis of differentially methylated regions. Briefly, genomic DNA was isolated from HL-7702 cell lines and 2 μg DNA was treated with sodium bisulfite using an EZ DNA Methylation-Gold Kit (ZYMO Research, USA) according to the manufacturer’s instructions. The specific primers based on the reverse complementary strands of the TFEB-promoter was designed using EpiDesigner software (Agena, USA), and the quantitative results for each CpG or multiple CpGs were analysed in EpiTYPER^™^ (Agena, USA). The primer sequences were 5’-aggaagagagAGGTATTTAAGGGTATTTTTGGTGG-3’ and 3’-cagtaatacgactcactataggg agaaggctCCTATAATCCCAACATTTTAAAAAACC-5’. Bisulfite-modified DNA PCR amplifications were performed with precycling hold at 94°C for 4 min and subjected to 45 cycles of 94°C for 20 seconds, 56°C for 20 seconds, and at 72°C for 1 min, and a final extension at 72°C for 3 min. Further experimental analysis of the contents of DNA methylation was determined, as described previously [18].

### mRFP-GFP-LC3 fluorescence microscopy

mRFP-GFP-LC3 adenoviral vectors were obtained from HanBio Technology Co. Ltd. (Shanghai, China). Hepatocyte autophagy was analysed using tandem mRFP-GFP-LC3 fluorescence microscopy. Hepatocytes were cultured in 35 mm confocal dishes for 24 h and then infected with mRFP-GFP-LC3 adenovirus for 2 h. Then, 100 μmol/L Hcy was added to the medium for 24 h at 37°C. Yellow and red puncta were observed using an Olympus BX51 confocal fluorescence microscope (Tokyo, Japan).

### Immunofluorescence

Hepatocytes were fixed with 4% paraformaldehyde for 20 min at room temperature (RT), permeabilized with 0.1% Triton on ice, and then blocked in PBS containing 5% bovine serum albumin (BSA) for 1 h, followed by incubation with a specific antibody against TFEB (ab267351, Abcam) overnight at 4°C. After that, Alexa Fluor™ conjugates (Life Technologies, USA) against the secondary antibody and DAPI were applied for 2 h at RT. Cells were then imaged by laser confocal microscopy (Olympus BX51, Japan).

### Chromatin immunoprecipitation (ChIP) assays

Formaldehyde at 1% was applied to the hepatocytes to cross-link for 15 min, and then glycine was replenished to a maximum concentration of 125 mM. Cells were flushed with cold PBS, collected, and then sonicated on a 20% power ultrasonic lyser to cut the DNA into fragments with an average size of 200 to 1000 bp. Then 50 μL from each sonicated sample was pipetted, and the fragment size and DNA concentration were measured. Cell lysates were incubated overnight with 30 μL ChIP-grade Protein G agarose beads (Merck Millipore, Germany) and 10 µg DNMT1, DNMT3a, and DNMT3b antibodies (Abcam, USA). Agarose beads were gathered and sequentially exposed to proteinase K at 60°C for 3 h and RNase at 37°C for 2 h. DNA purification was performed using a SpinPrep™ PCR Clean-up Kit (Merck Millipore, Germany). DNA fragments were assayed by qRT-PCR using the primer sequences listed below: forwards primer: 5’-CTGGTATTAGCCAGAACATGTCAG-3’, reverse primer: 5’-CCTCTTGCACAGTATGTAGCACC-3’. Samples were standardized to input DNA.

### Statistical analysis

The Results are presented as the means ± standard deviations (SD) and statistical significance was analysed with one-way analysis of variance (ANOVA), followed by the Newman–Keuls test. Each test was carried out three times.

## RESULTS

### Hcy promotes autophagy in hepatocytes

We used 100 μmol/L Hcy to stimulate hepatocytes, to verify whether Hcy induces hepatocyte autophagy. We examined the conversion of LC3BI to LC3BII (a marker for autophagy activation) and p62 expression (a marker of autophagy inhibition), as well as the initiation of macroautophagy flux. The results showed that Hcy increased the ratio of LC3BII/I and inhibited p62 expression in hepatocytes (Fig. 1A and B). Interestingly, hepatocytes were infected with a pH-sensitive tandem mRFP-GFP-LC3 adenovirus to monitor autophagy-induced puncta formation. Yellow puncta reflect the combination of mRFP and GFP fluorescence labels autophagosomes, whereas sole red puncta (mRFP only) labels autolysosomes due to the quenching of GFP fluorescence under acidic pH conditions. As shown in Fig. 1C, free red and yellow puncta significantly increased in hepatocytes, indicating an increase in both autophagosomes and autolysosomes. Collectively, these data indicated that Hcy induced autophagy in hepatocytes.

**Fig. 1.**
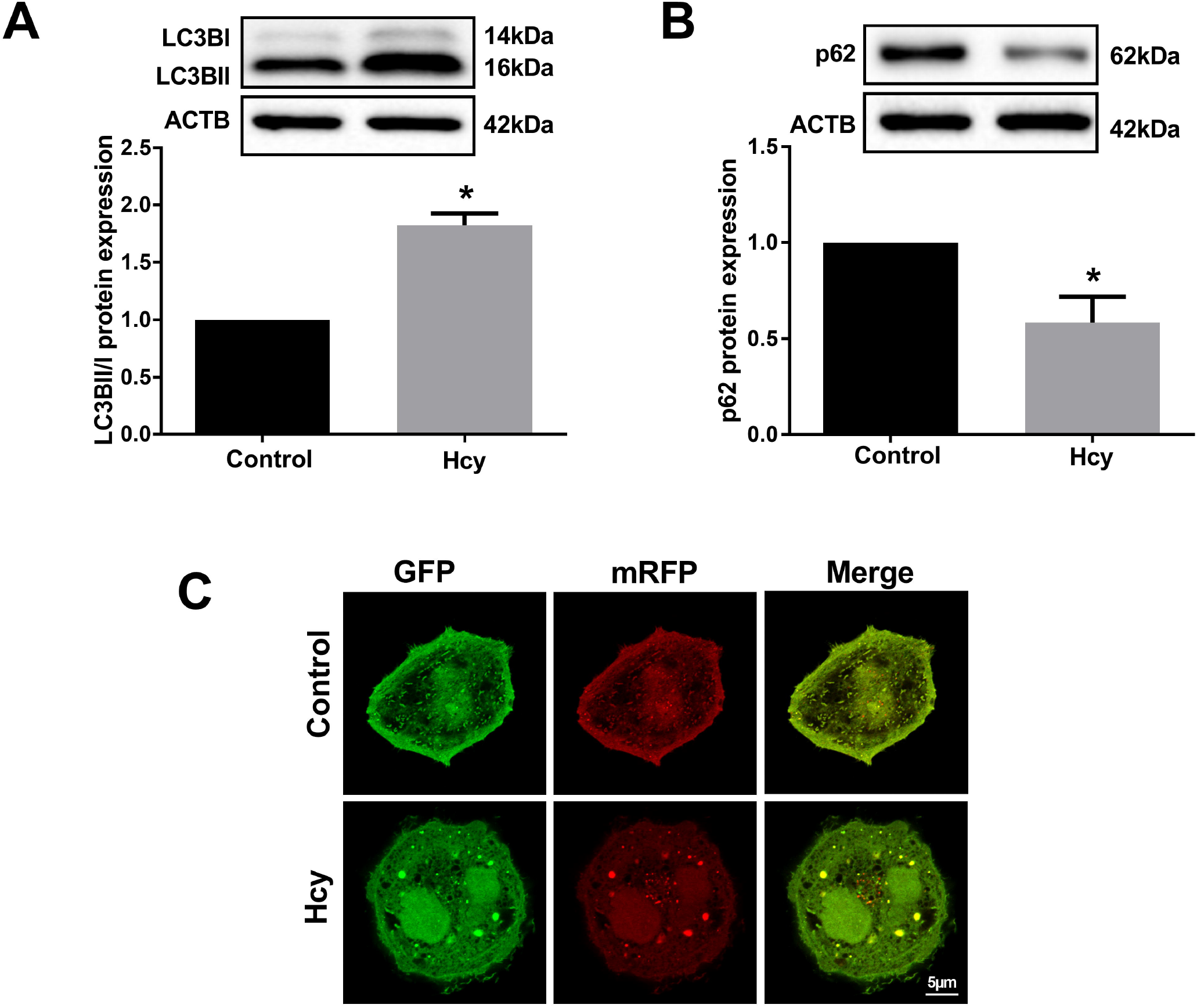
The autophagy assay in Hcy-induced hepatocytes. (A, B) The protein levels of LC3BII/I and p62 in Hcy-induced hepatocytes were measured by immunoblot (Hcy: 100 μmol/L). (C) Tandem mRFP-GFP-LC3 fluorescence microscopy was performed to analyze hepatocyte autophagy. Liver cells were incubated with Hcy (100 μmol/L) for 24 h (scale bars=5 μm). Yellow puncta, the merge of mRFP and GFP fluorescence, indicate autophagosomes, whereas free red puncta (mRFP only) represent autolysosomes. Data represent the mean ± SD from three independent experiments, \**P* < 0.05.

### TFEB plays an essential role in the promoting of hepatocyte autophagy induced by Hcy

To determine whether TFEB is involved in Hcy-induced liver autophagy, TFEB mRNA and protein expression were analysed by qRT-PCR and immunobloting in hepatocytes exposed to Hcy. We found that TFEB expression increased in hepatic cells exposed to Hcy (*P*<0.05, Fig. 2A). Additionally, to verify the distribution of TFEB in hepatocytes, laser confocal microscopy showed that TFEB was primarily distributed in the nucleus. The fluorescence signal intensity in hepatocytes exposed to Hcy was stronger than that in the control, indicating that Hcy promoted the expression of TFEB (Fig. 2B). Then, TFEB shRNA adenovirus (sh-TFEB) was infected into hepatocytes to verify its interference efficiency (Fig. 2C). We found that knockdown of TFEB expression inhibited the proportions of LCB3II/I and enhanced the expression of p62 in hepatocytes (Fig.2D). The results indicate that TFEB plays an important role in Hcy-induced autophagy in hepatocytes.

**Fig. 2.**
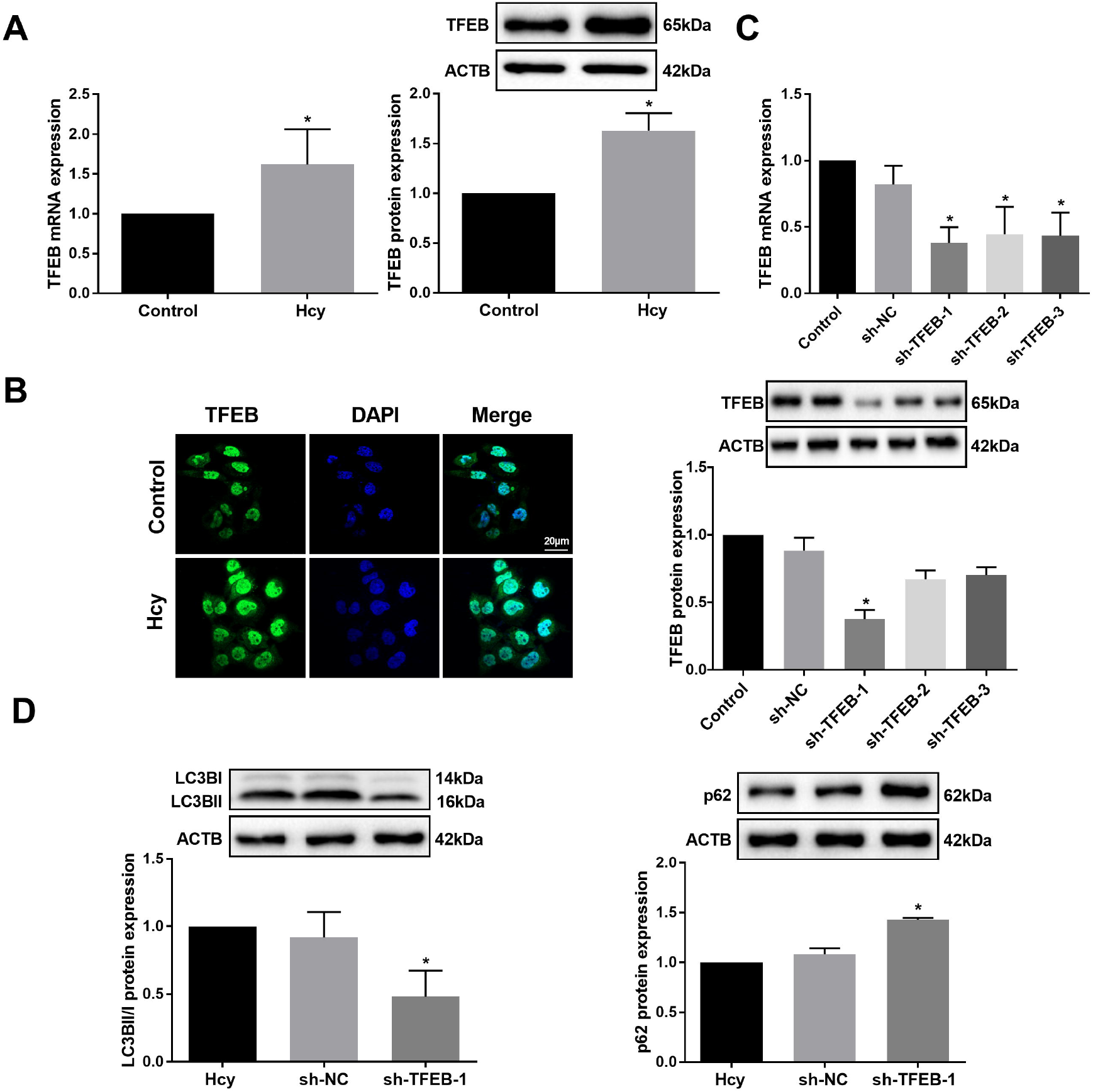
Hcy promotes autophagy by upregulating TFEB levels in hepatocytes. (A) TFEB mRNA and protein expression in hepatocytes were measured by qRTLPCR and immunobloting, respectively. (B) Immunofluorescent staining of TFEB (green) in hepatocytes after treatment with Hcy, Nuclei were stained with DAPI (blue) (scale bars=20 μm). (C) mRNA and protein levels of TFEB were measured by qRTLPCR and immunobloting in hepatocytes after infection with sh-TFEB adenovirus (sh-TFEB-1, sh-TFEB-2, sh-TFEB-3) or sh-NC (control). (D) The protein expression of LC3BII/I and p62 was detected by immunoblot in Hcy-induced hepatocytes infected with sh-TFEB-1 adenovirus. Data represent the mean ± SD from three independent experiments, \**P* < 0.05.

### TFEB hypomethylation plays an essential role in upregulating TFEB expression in hepatocytes

To investigate whether Hcy modulates TFEB expression via CpG methylation, we used MethPrimer software to predict CpG island changes (+483 bp/+953 bp) in the TFEB promoter region. The Methylation status of CpG islands in the TFEB promoter was measured by nMS-PCR and MassARRAY (Fig. 3B, C). As expected, Hcy reduced DNA methylation of TFEB promoter in hepatocytes, and 5-azacytidine (AZC) further inhibited this process. To further understand the effects of DNA methylation on TFEB expression, we used AZC to intervene in Hcy-induced hepatocytes and found increased TFEB mRNA and protein levels (Fig.3D), suggesting that Hcy downregulates the TFEB DNA methylation and upregulates TFEB expression in hepatocytes.

**Fig. 3.**
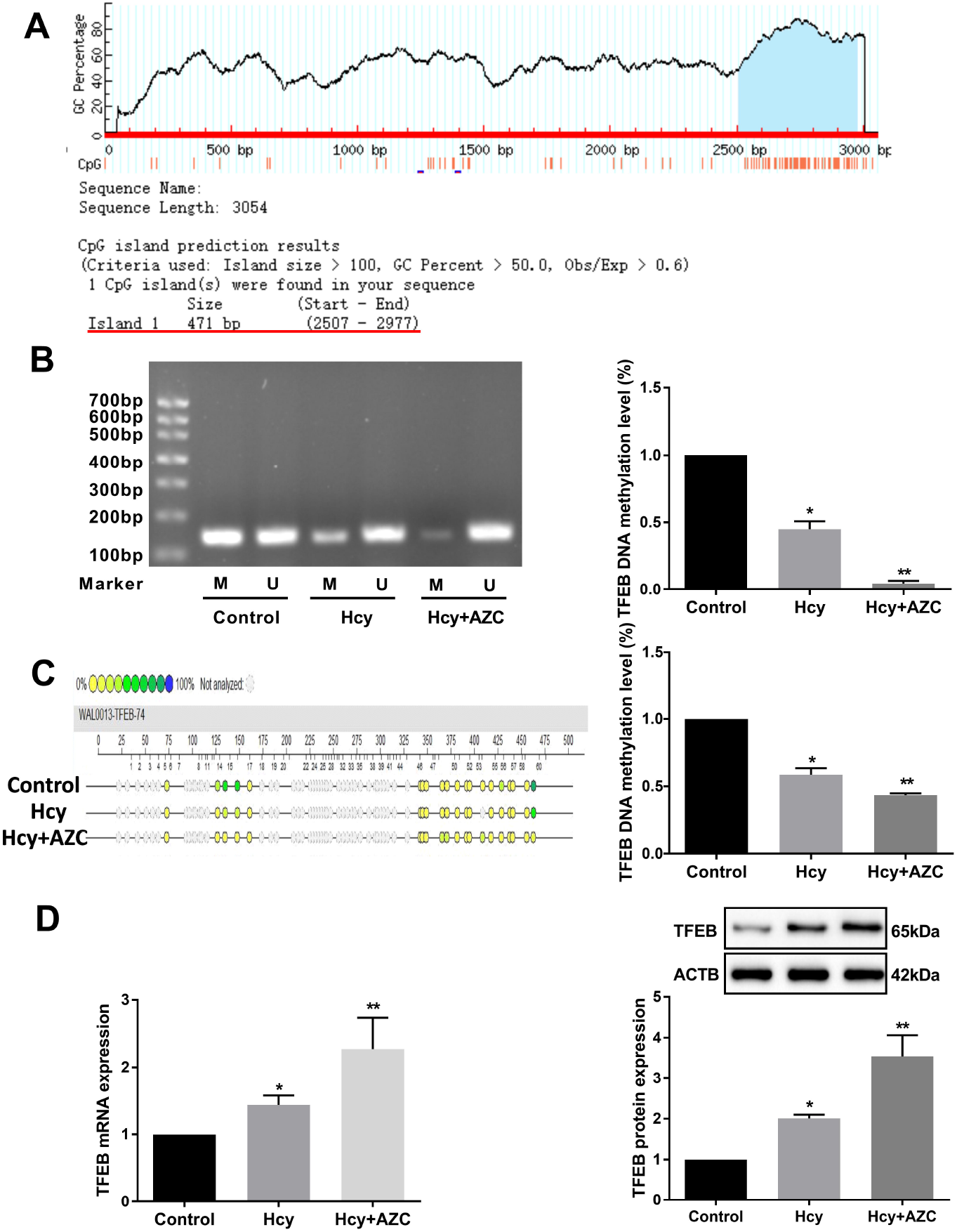
Hcy downregulates the level of TFEB DNA methylation. (A) Screenshots of the putative CpG islands in the 5’-flank regions of the TFEB gene. (B) The level of DNA methylation in the TFEB promoter was detected by nMS-PCR in hepatocytes after exposure to Hcy or Hcy plus 5-azacytidine (AZC). M indicates the methylated PCR band; U indicates the unmethylated PCR band. (C) MassARRAY was used to examine the methylation status of TFEB promoter in hepatocytes. The circle color represents the percentage of methylation in each CpG site. Blue indicates methylation (100%) and yellow indicats a lack of methylation (0%). Gray circles represent the unanalysed CpG sites. (D) The mRNA and protein expression of TFEB was measured by qRT-PCR and immunobloting in hepatocytes exposed to Hcy and AZC. Data represent the mean ± SD from three independent experiments, \**P* < 0.05, \*\**P* < 0.01.

### DNMT3b positively modulates Hcy-induced TFEB DNA methylation in hepatocytes

To understand the role of DNMTs in regulating TFEB DNA methylation in hepatocytes, cells were exposed to DC-05, TF-3, and Nanaomycin A (NA), which are DNMT1, DNMT3a and DNMT3b inhibitors, respectively. Interestingly, NA significantly increased TFEB expression in hepatocytes compared to other inhibitors (Fig.4A). ChIP further demonstrated that DNMT3b, but not DNMT1 or DNMT3a, bound less to the proximal promoter region of TFEB after exposure to Hcy than in the control group in hepatocytes (Fig.4B).

To gain insight into the role of DNMT3b in the regulation of DNA methylation in the TFEB promoter, we infected DNMT3b overexpressing adenovirus (Ad-DNMT3b) into hepatocytes and detected the expression of DNMT3b. The data showed that DNMT3b mRNA and protein levels were increased (Fig.4C). Meanwhile, we screened the most functional DNMT3b interference adenovirus (sh-DNMT3b) and verified its efficiency. Remarkably, sh-DNMT3b-1 showed the best interference efficiency (Fig. 4D). Overexpression of DNMT3b upregulated TFEB DNA methylation and inhibited its RNA and protein expression, while knockdown of DNMT3b further inhibited TFEB DNA methylation in Hcy-induced hepatocytes (Fig. 4E-F). In addition, overexpression of DNMT3b increased its binding efficacy in the TFEB promoter and reduced autophagy in hepatocytes (Fig. 4G-H). Finally, using mRFP-GFP-LC3 adenovirus, we found that autophagosomes and autophagic lysosomes decreased in DNMT3b-overexpressing hepatocytes exposed to Hcy (Fig. 4I). Taken together, DNMT3b is a specific methyltransferase that regulates TFEB methylation and autophagy in Hcy-induced hepatocytes.

**Fig. 4.**
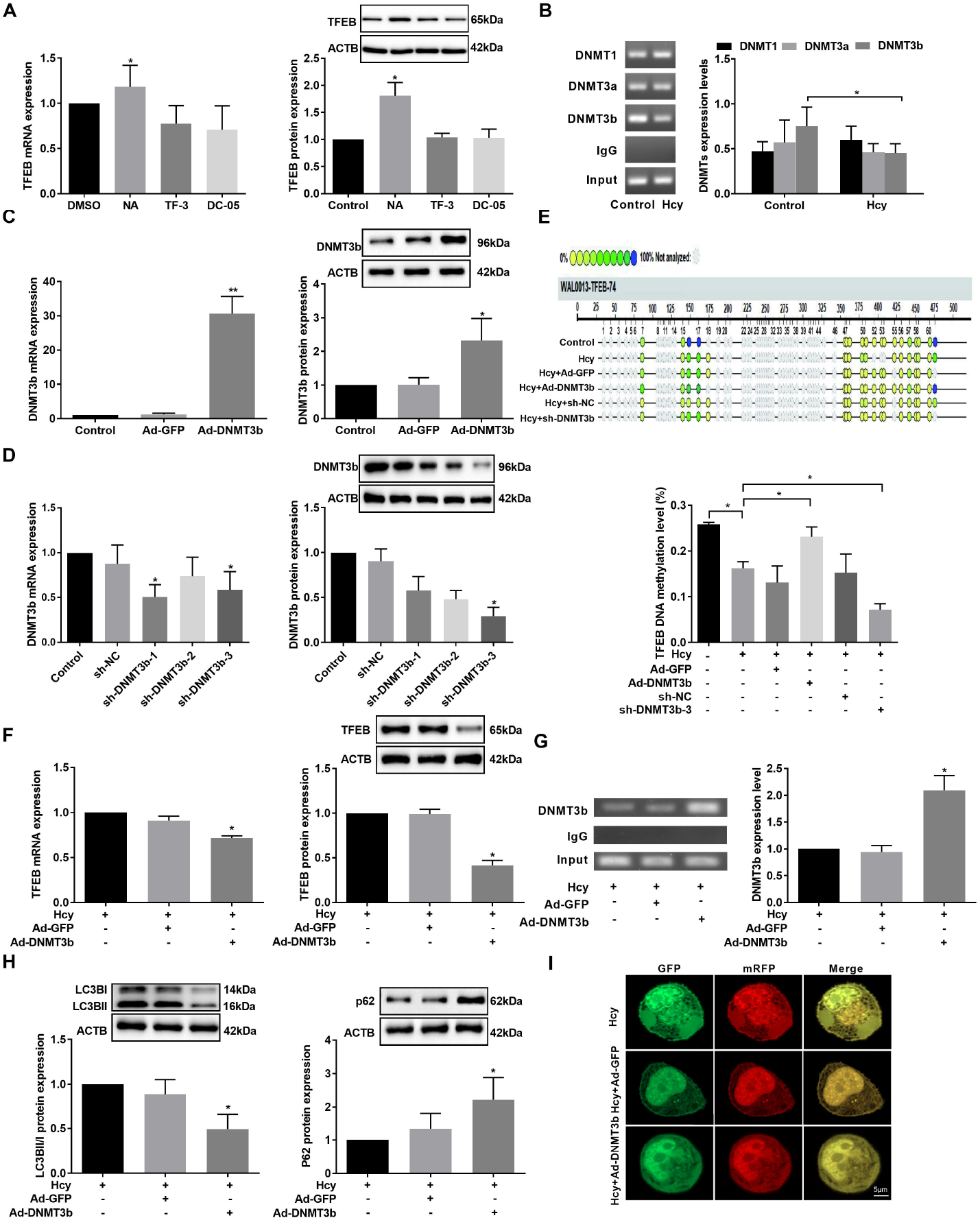
The effect of DNMT3b on the TFEB DNA methylation and autophagy. (A) The difference of TFEB mRNA and protein levels in hepatocytes exposed to Hcy and DC-05, TF-3, and Nanaomycin A (NA), the corresponded inhibitors of DNMT1, DNMT3a and DNMT3b, respectively. (B) The binding level between DNMT and TFEB promoter in hepatocytes was examined by ChIP assay. (C) DNMT3b mRNA and protein were measured by qRT-PCR and immunoblot in hepatocytes with DNMT3b adenovirus infection. (D) DNMT3b levels were measured by qRT-PCR and immunoblotting in hepatocytes infected with sh-DNMT3b adenovirus (sh-DNMT3b-1, sh-DNMT3b-2, sh-DNMT3b-3). (E) Quantifying DNA methylation of the TFEB promoter in hepatocytes after DNMT3b overexpression or inhibition. (F) The effect of DNMT3b overexpression on the TFEB levels in hepatocytes. (G) The binding level of DNMT3b to the TFEB promoter was measured by ChIP assay in hepatocytes exposed to Ad-DNMT3b. (H) Detection of LC3BII/I and p62 protein levels after overexpression of DNMT3b. (I) Confocal images of mRFP-GFP-LC3 fluorescent puncta in hepatocytes after being exposure to Ad-DNMT3b (scale bars = 5 μm). Data represent the mean ± SD from three independent experiments, \**P* < 0.05, \*\**P* < 0.01.

## DISCUSSION

In the current study, we examined the role of TFEB in Hcy-induced hepatocytes autophagy. Our findings confirmed the role of DNA hypomethylation in upregulating TFEB expression, which leads to Hcy-induced autophagy in hepatocytes.

Hcy is a sulfur-containing amino acid produced in internal metabolism [19]. As liver is one of the principal organs of Hcy metabolism, once liver function is impaired, it causes abnormal methionine metabolism, which leads to the release of accumulated Hcy into the plasma. Hcy, in turn, can affect liver function. Hcy has been linked to the pathogenesis of several diseases, including coronary artery disease and liver disease, both of which are characterized by elevated levels of total Hcy in plasma [20]. In the early stage, the HHcy model of ApoE^-/-^ mice was replicated by feeding with a high methionine diet, and serum Hcy was detected to verify model establishment [21]. We also found that HHcy caused liver damage in ApoE^-/-^ mice, and Hcy increased the level of autophagy in the liver tissue of ApoE^-/-^ mice.

TFEB is a member of the leucine zipper family of transcription factors. It was found that TFEB nuclear translocation increased transcription of genes encoding autophagic and lysosomal proteins, thereby promoting lysosomal biogenesis and autophagosome formation, and increasing autophagy [22]. Moreover, ethanol activation of mTORC1 disrupted TFEB-mediated liver lysosomal biogenesis, resulting in autophagy deficiency in mice. In contrast, the overexpression of TFEB increases the biogenesis of lysosomes and prevents ethanol-induced steatosis and liver injury [23]. These studies suggest that liver autophagy and lysosome function can be regulated by changing the activity of TFEB. We found that increased TFEB boosted autophagy in Hcy-induced hepatocytes. Conversely, knockdown of TFEB decreased this effect.

Elevated Hcy regulates genomic DNA methylation and induces specific methylation in the promoter regions of genes associated with human diseases, including carcinoma, mental illness, neurodegenerative disease, and cardiovascular disease [24-26]. DNA methylation is a biological process in which a methyl group is covalently linked with cytosine to produce 5-methylcytosine (5mC) [27]. DNA methylation has been widely considered an epigenetic silencing approach that plays a role in numerous cellular metabolic events, including X-chromosome inactivation, genomic imprinting, and gene transcription [28]. The methylation process is catalyzed by a group of enzymes called DNMTs. DNMT3a and DNMT3b are responsible for the de novo methylation mode, whereas DNMT1 is responsible for maintaining methylation. Therefore, Hcy is an important intermediate that plays a crucial role in DNA methylation [29].

Numerous studies have demonstrated that methylated sequences are not recognized by transcription factors, which prevents the expression of corresponding genes. DNA methylation of cytosine residues transforming to 5-methylcytosine may assist in this process, so the regulatory regions of these genes are frequently hypomethylated, which results in gene overexpression, and hypermethylated in the cells and tissues, which silences these genes [30]. The methyl group used for methylation reactions originates from S-adenosylmethionine (SAM), an intermediate in the metabolism of Hcy. Upon methylation, SAM is converted to S-adenosylhomocysteine (SAH), which inhibits transmethylation reactions. HHcy could lead to global hypomethylation through SAH aggregation and weakened methylation capacity (decreased SAM/SAH ratio) [31]. Ma *et al*. found Hcy induced the extracellular-superoxide dismutase (EC-SOD) DNA hypomethylation in macrophages and that DNMT1 acted as the essential enzyme in the methyl transfer process leading to downregulation of EC-SOD expression and increased atherosclerosis [32]. In this study, we found that the level of TFEB DNA methylation is decreased in Hcy-incubated hepatocytes. Then, Hcy mainly affected DNMT3b to upregulate TFEB expression. Using an adenovirus overexpressing with DNMT3b, we found that the methylation levels of TFEB were elevated, the expression of TFEB was decreased, and the level of hepatocyte autophagy also suppressed.

In conclusion, our data reveal that TFEB plays a key role in Hcy-induced hepatocyte autophagy. Meanwhile, TFEB DNA hypomethylation upregulated TFEB expression, which is a new mechanism by which Hcy promotes hepatocyte autophagy (Fig. 5). Nevertheless, the potential role of TFEB in Hcy-induced autophagy needs to be further investigated.

**Fig. 5.**
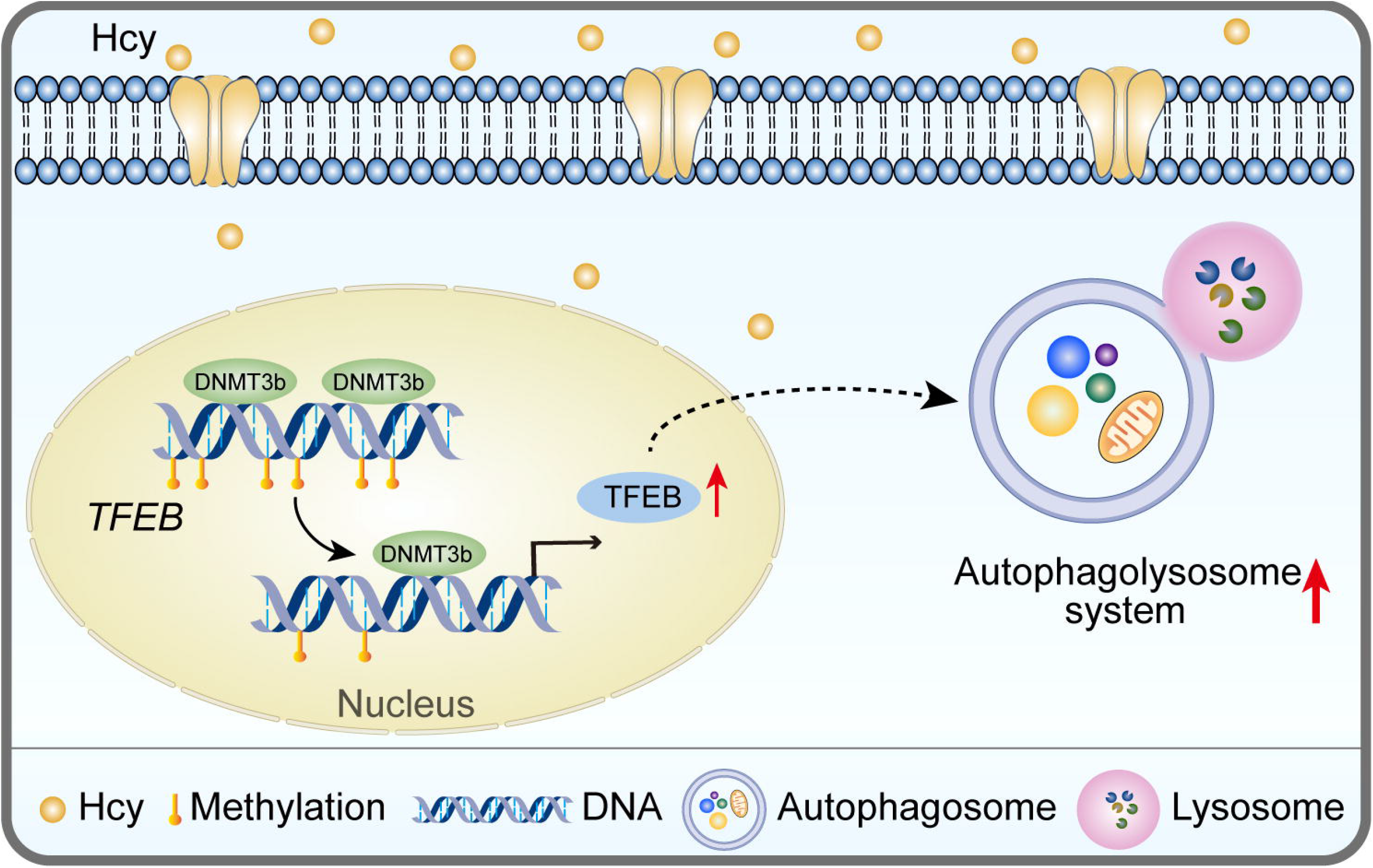
Homocysteine accelerates hepatocyte autophagy by upregulating TFEB via DNMT3b-mediated DNA hypomethylation.

## Acknowledgements

This study was supported by grants from the National Natural Science Foundation of China (U21A20343, 82160088, 81870332, 81700404, 82260088, and 82060110); Natural Science Foundation of Ningxia Province (2022AAC05025, 2020AAC02021, 2020AAC02038 and 2021AAC03337); Key Research and Development Projects in Ningxia Province (2019BFG02004, 2020BFH02003, 2020BEG03005 and 2020BEG03008); Ningxia Project of Supporting Engineering Talents (TJGC2019092); A School-level special Talents launching project of Ningxia Medical University (XT2018015); Ningxia Science and Technology Leading Talent Project (KJT2017007); and Basic scientific research operating expenses from the public welfare research institutes at the central level of the Chinese Academy of Medical Sciences (2019PT330002).

## Author’ contributions

Anning Yang, Wen Zeng, Yinju Hao, and Hongwen Zhang contributed to the concept of the study, they also analysed and wrote the manuscript, Qingqing Wang, Ning Ding, Shangkun Quan, and Yue Sun helped to perform experiments and acquire data, Jianmin Sun, Xiaoling Yang and Huiping Zhang mainly provided some help for our investigation. Yideng Jiang, Yun Jiao, Kai Wu and Bin Liu designed and supervised the study.

